## Supplementary figures and images for "PI(3,4,5)P3 allosteric regulation of repressor activator protein 1 controls antigenic variation in trypanosomes"

### Figure 1-figure supplement 1

Figure 1 – figure supplement 1

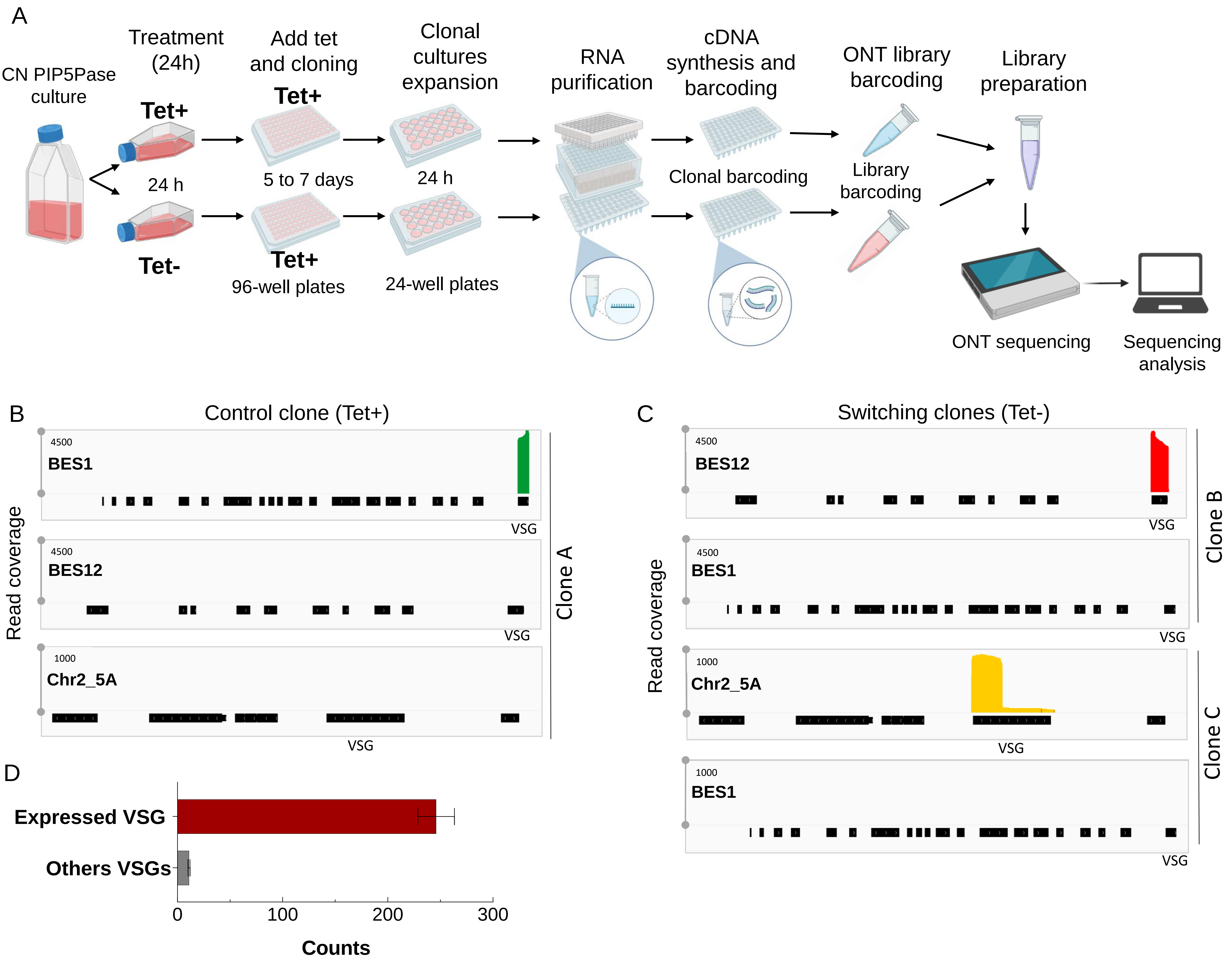

### Figure 1-figure supplement 2

Figure 1 – figure supplement 2

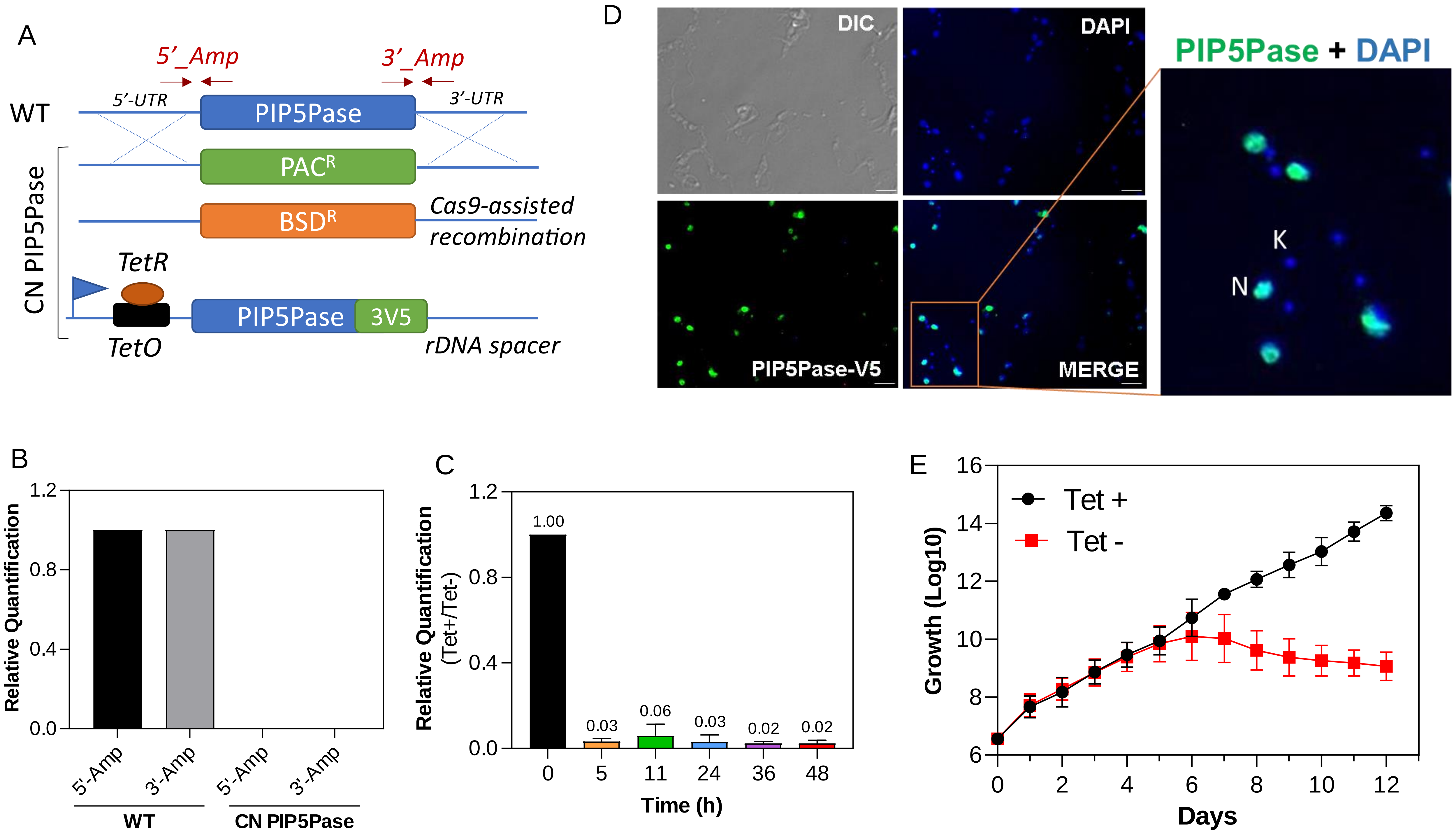

### Figure 2-figure supplement 1

Figure 2 – figure supplement 1

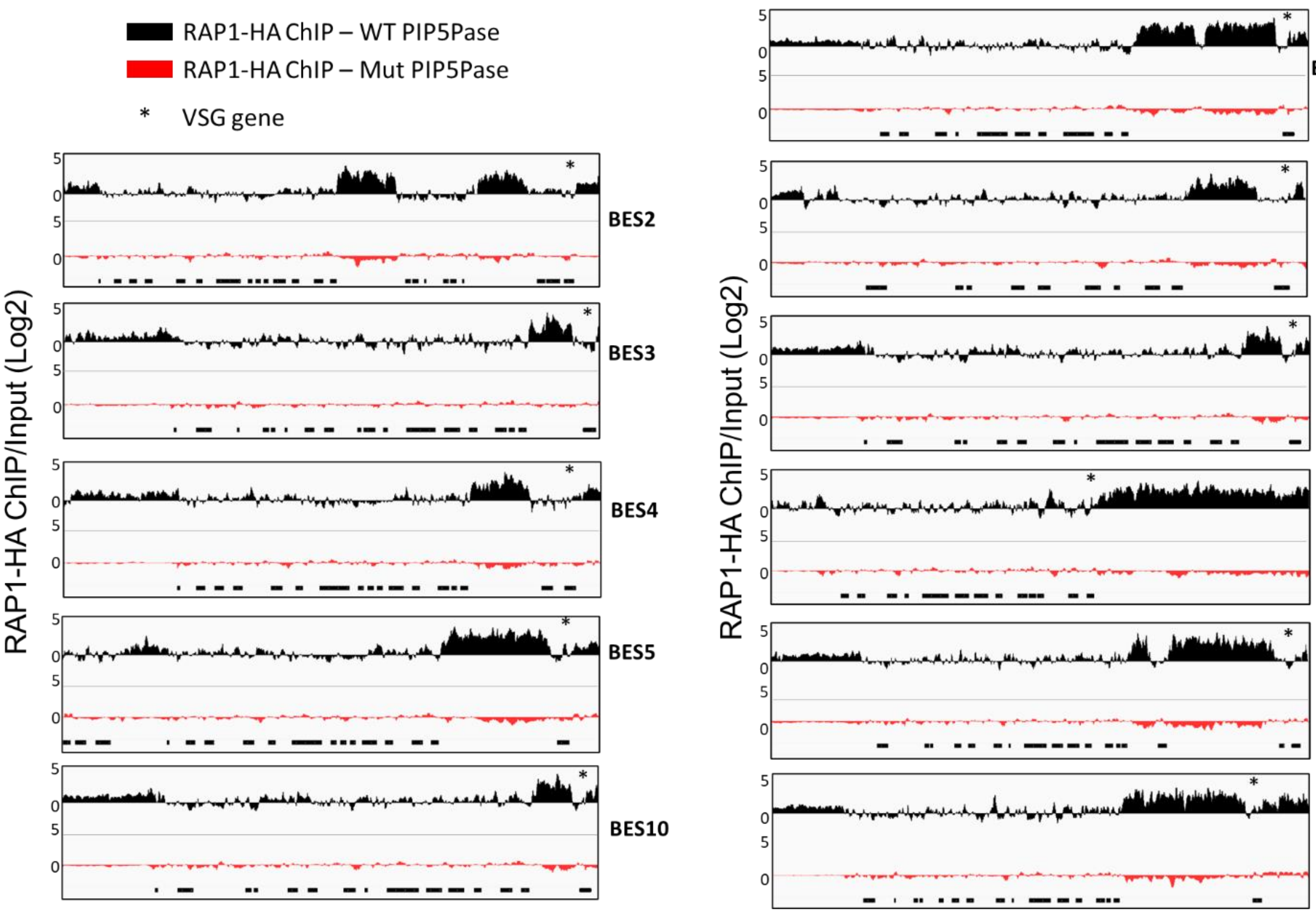

### Figure 2-figure supplement 2

Figure 2 – figure supplement 2

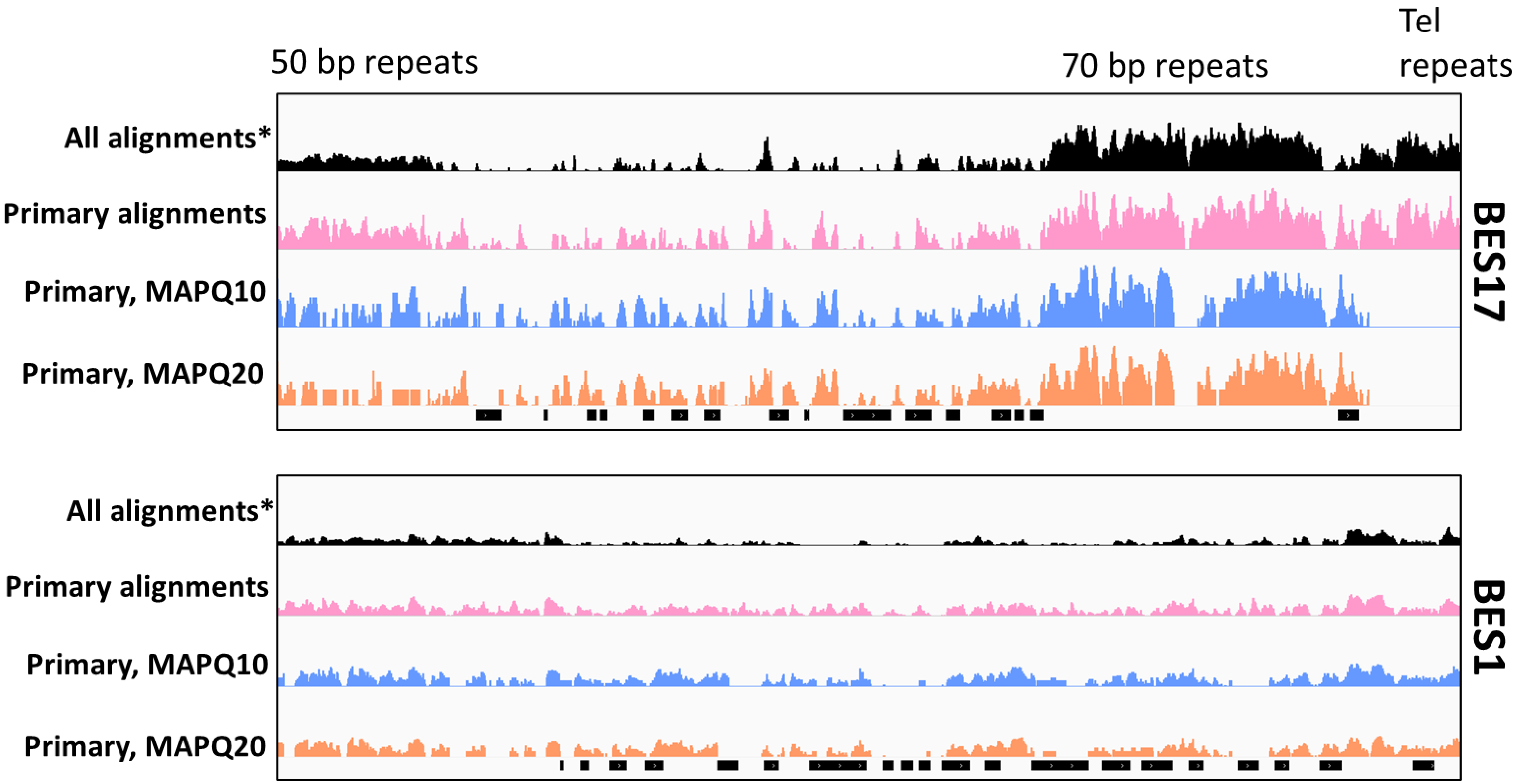

### Figure 2-figure supplement 3

Figure 2 – figure supplement 3

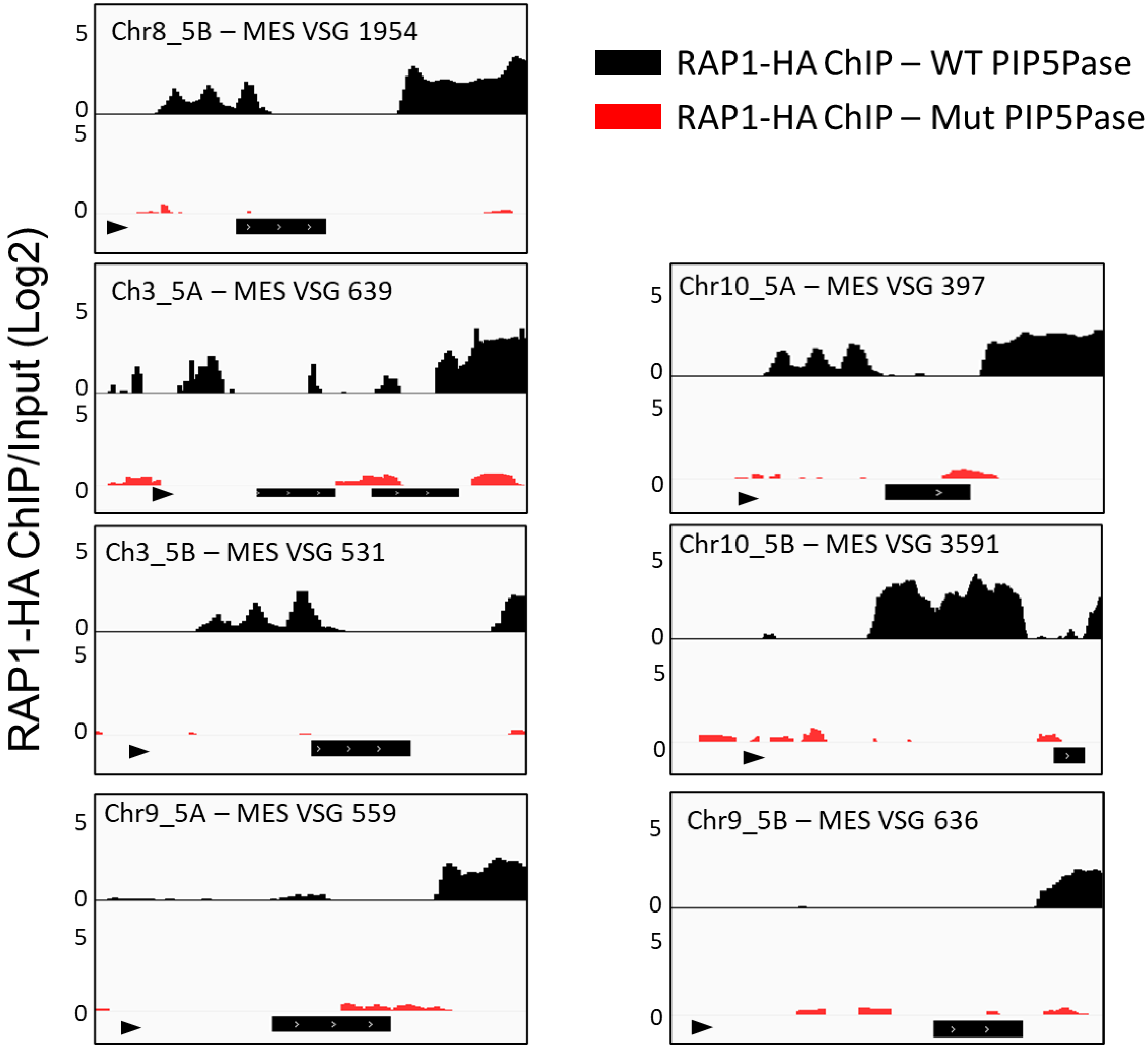

### Figure 3-figure supplement 1

Figure 3 – figure supplement 1

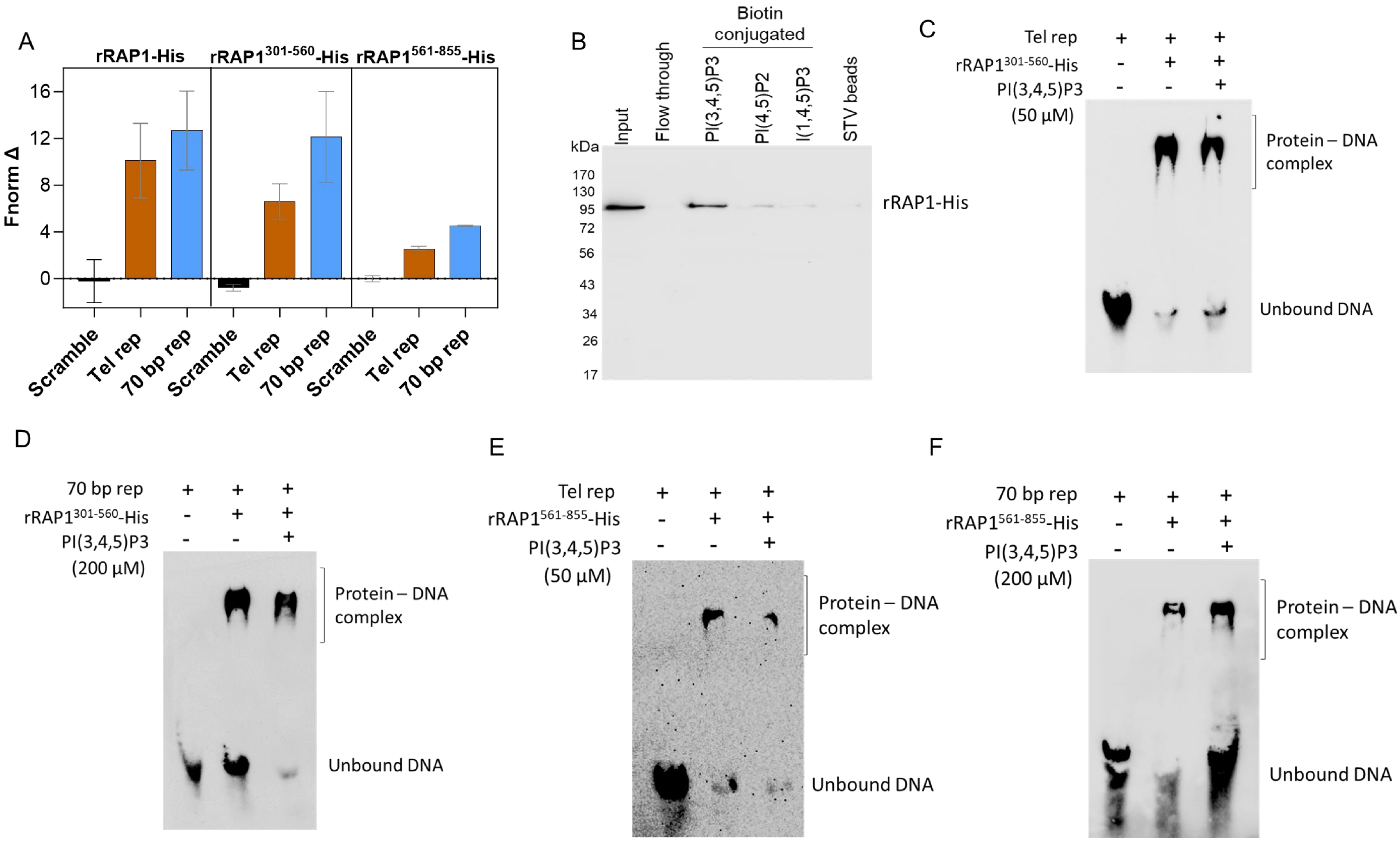

### Figure 4-figure supplement 1

Figure 4 – figure supplement 1

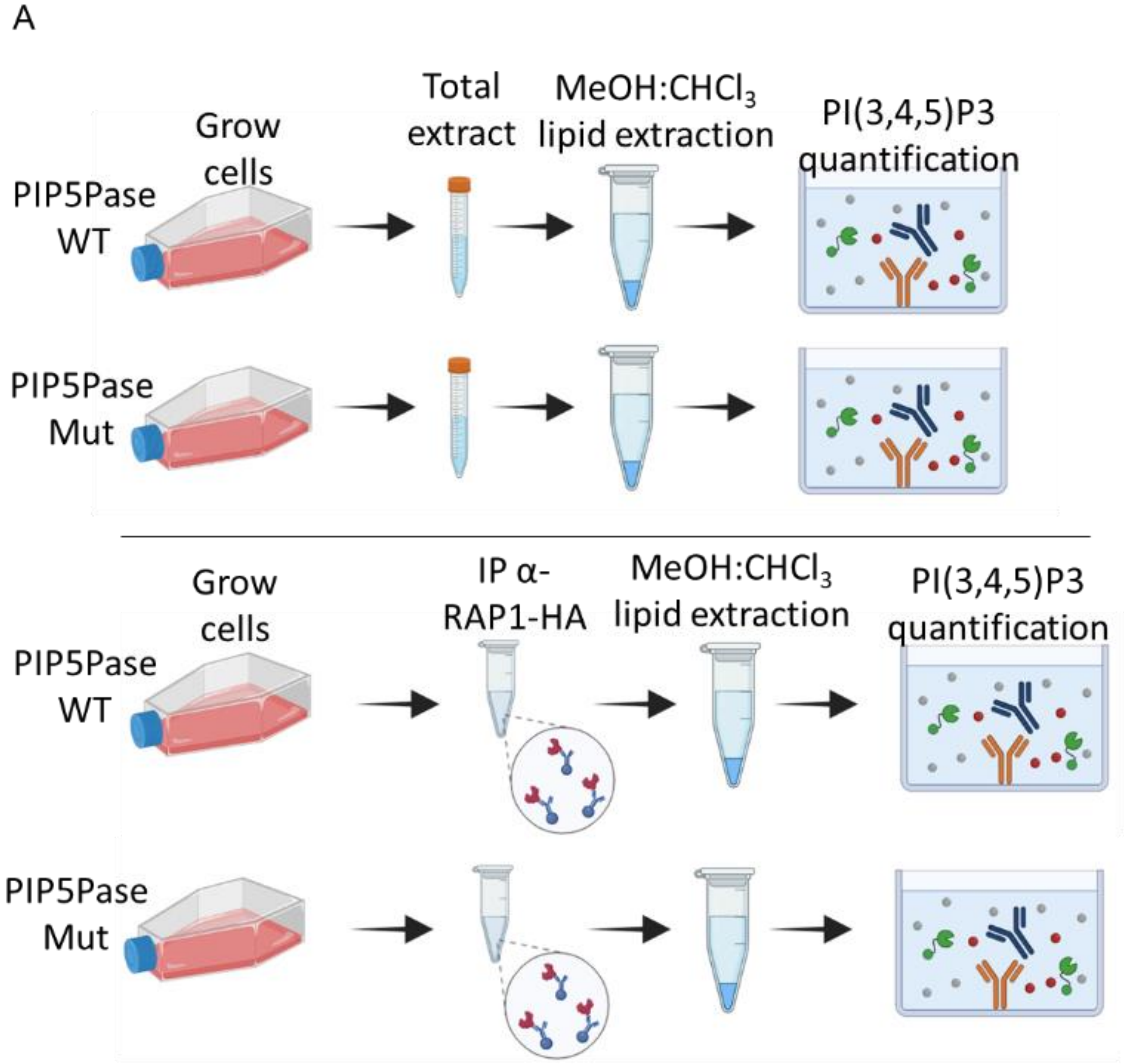


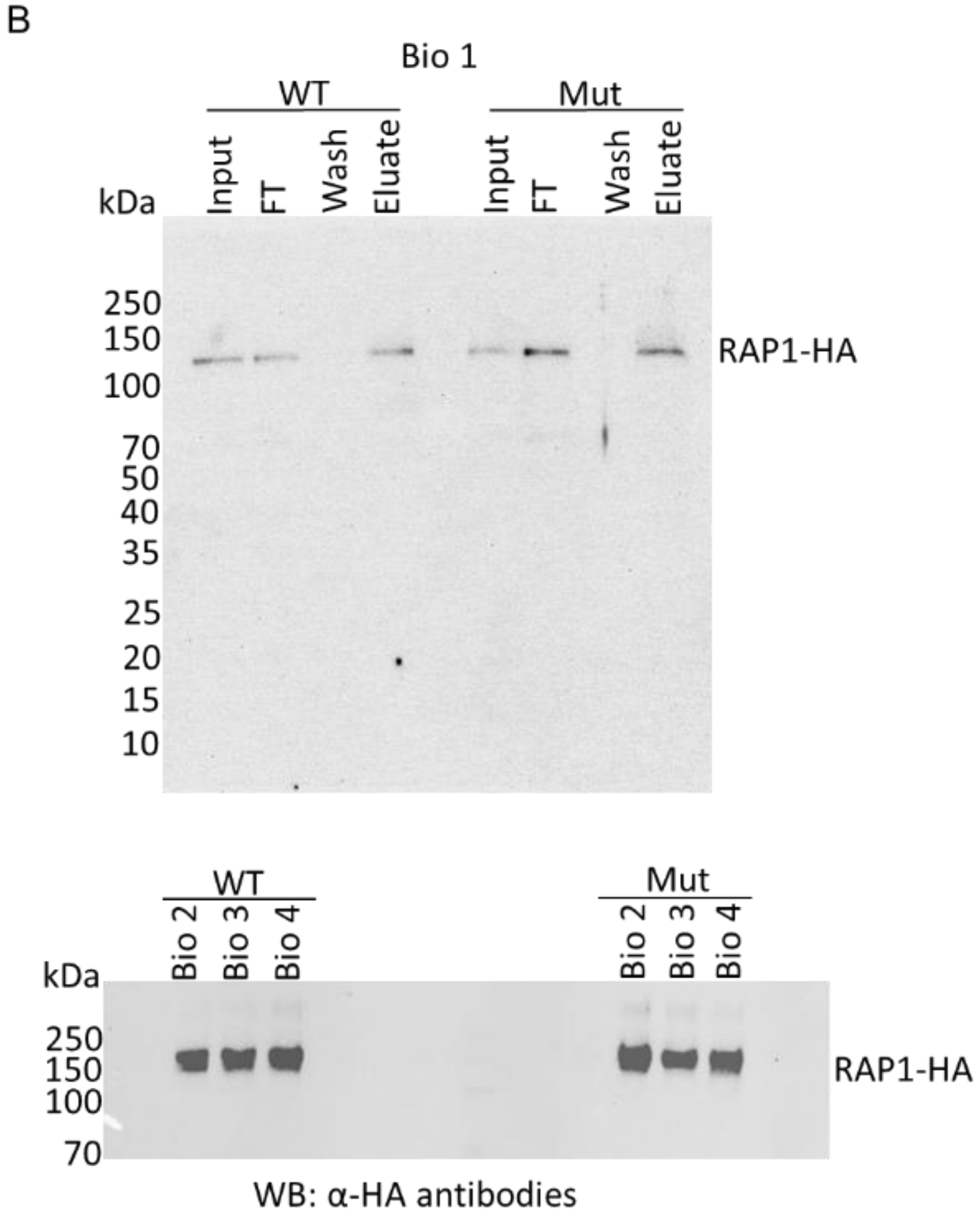

### Figure 5-figure supplement 1

Figure 5 – figure supplement 1

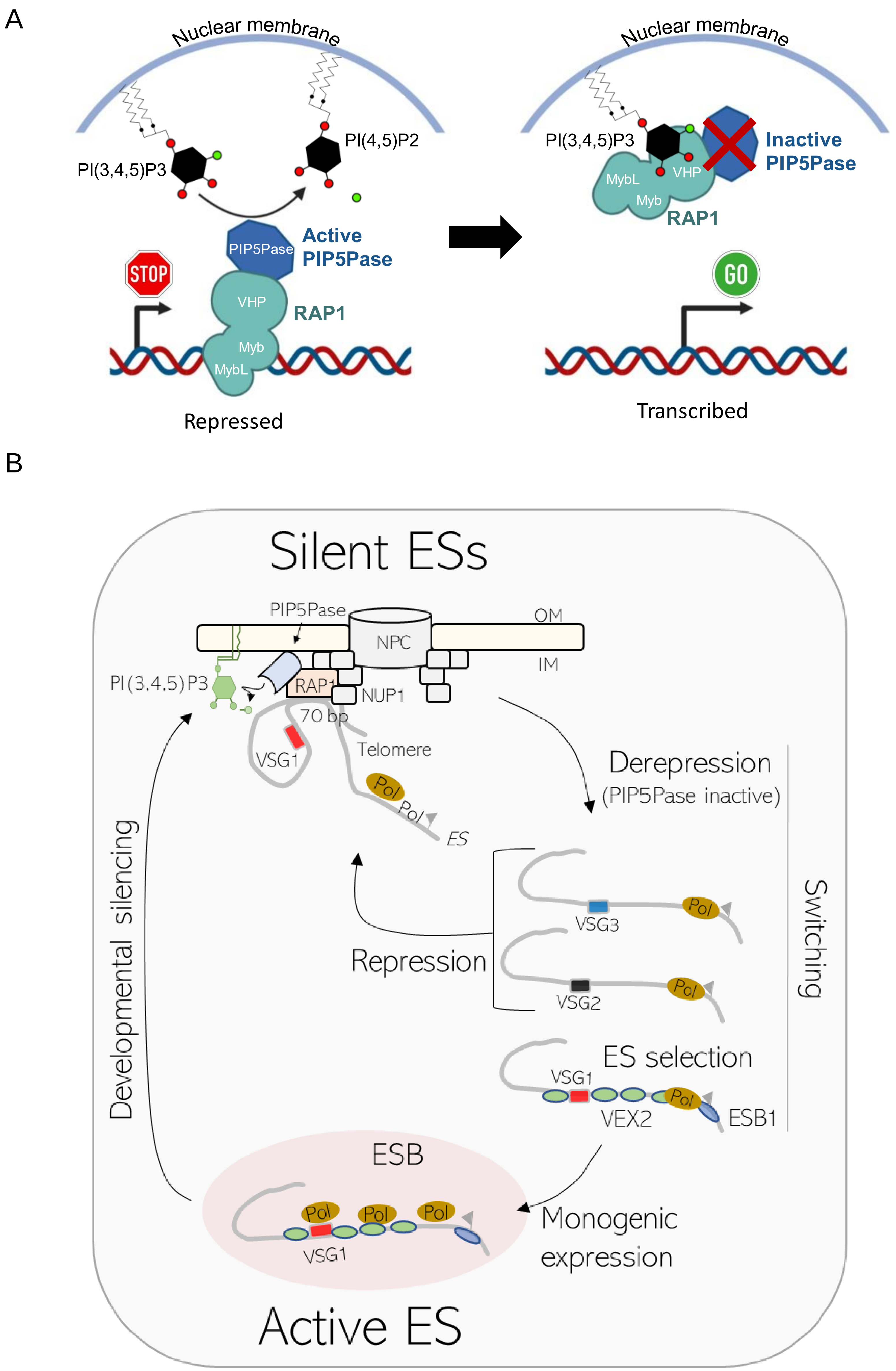
