## Supplementary File 3 for "PI(3,4,5)P3 allosteric regulation of repressor activator protein 1 controls antigenic variation in trypanosomes"

| **Chr** | **GeneID** | **WT-Bio1** | **WT-Bio2** | **WT-Bio3** | **Mut-Bio1** | **Mut-Bio2** | **Mut-Bio3** | **Short name** | **Description** |
| --- | --- | --- | --- | --- | --- | --- | --- | --- | --- |
| BES1 | Tb427_000016000 | 19.88 | 18.31 | 17.26 | 15.63 | 15.13 | 16.78 | BES1_VSG2 | Trypanosome variant surface glycoprotein (A-type), putative |
| Chr1_3A | Tb427_000240100 |  |  |  | 16.14 | 16.11 | 15.92 | Chr1_3A_VSG | Trypanosomal VSG domain containing protein, putative |
| unitig_23 | Tb427_000631200 |  |  |  | 9.02 | 8.51 | 9.24 | Unitig_VSG-23 | Tb427VSG-23_unitig_Tb427v10 |
| BES12 | Tb427_000008000 | 7.82 |  |  | 8.07 | 7.57 | 8.69 | BES12_VSG | Trypanosome variant surface glycoprotein (A-type) |
| Chr8_5A | Tb427_000505400 |  |  |  | 7.07 | 8.15 | 8.57 | Chr8_5A_VSG-2261 | VSG Tb427VSG-2261 |
| Chr8_5A | Tb427_000516000 |  |  | 5.94 | 9.02 | 7.31 | 8.43 | Chr8_5A_VSG | Trypanosomal VSG domain containing protein, putative |
| Chr11_5A | Tb427_000173600 |  | 3.83 | 3.94 | 7.98 | 8.38 | 8.02 | Chr11_5A_VSG | Trypanosome variant surface glycoprotein (A-type), putative |
| Chr5_3A | Tb427_000349200 |  |  | 6.53 | 7.66 | 7.44 | 7.92 | Chr5_3A_VSG | Trypanosomal VSG domain containing protein, putative |
| BES15 | Tb427_000012400 | 3.91 | 3.83 |  | 6.66 | 6.31 | 7.43 | BES15_VSG | Trypanosomal VSG C-terminal domain containing protein, putative |
| BES7 | Tb427_000025600 |  |  |  |  | 4.98 | 6.69 | BES7_VEP | Trypanosomal VSG C-terminal domain containing protein, putative |
| Chr10_3B | Tb427_000080300 |  |  |  | 4.07 | 3.98 | 6.11 | Chr10_3B_VSG | Trypanosomal VSG domain containing protein, putative |
| Chr1_3A | Tb427_000236500 |  |  |  | 4.07 | 4.98 | 6.11 | Chr1_3A_VSG-699 | 100% identify to VSG Tb427VSG-699 |
| unitig_219 | Tb427_000732900 |  | 3.83 | 5.53 | 6.07 | 6.57 | 5.69 | Unitig_VSG | 96% identity to VSG 221, VSG expression site BAC |
| Chr8_5A | Tb427_000517200 |  |  |  | 5.07 | 6.31 | 5.69 | Chr8_5A_VSG-2088 | 100% identity to VSG Tb427VSG-2088 |
| Chr9_3A | Tb427_000555300 |  |  | 3.94 | 6.07 | 4.98 | 5.11 | Chr9_3A_VEP | Trypanosomal VSG C-terminal domain containing protein, putative |
| Chr8_3A | Tb427_000494500 |  |  |  | 6.39 | 5.57 | 4.11 | Chr8_3A_VSG-457 | 99% similar to Trypanosoma brucei VSG 547 gene |
