## Supplementary File 4 for "PI(3,4,5)P3 allosteric regulation of repressor activator protein 1 controls antigenic variation in trypanosomes"

| **Chromosome** | **Start** | **End** | **Length** | **Pileup** | **Location** | **FE** | ***p*-value -log10** | ***q*-value -log10** |
| --- | --- | --- | --- | --- | --- | --- | --- | --- |
| BES2 | 37 | 4971 | 4935 | 878.36 | 50 bp repeats | 1.27 | 11.88 | 10.19 |
| BES2 | 42238 | 50979 | 8742 | 710.97 | 70 bp repeats upstream VSG gene | 2.73 | 119.08 | 116.56 |
| BES2 | 63552 | 70309 | 6758 | 725.77 | 70 bp repeats upstream VSG gene | 2.49 | 102.10 | 99.74 |
| BES2 | 71419 | 75797 | 4379 | 886.5 | 70 bp repeats upstream VSG gene | 1.15 | 4.64 | 3.36 |
| BES2 | 79041 | 82168 | 3128 | 923.94 | downstream VSG gene and telomeric repeats | 1.23 | 9.32 | 7.66 |
| BES3 | 404 | 12982 | 12579 | 891.29 | 50 bp repeats | 1.19 | 7.07 | 5.58 |
| BES3 | 56564 | 61552 | 4989 | 647.06 | 50 bp repeats | 2.49 | 93.53 | 91.34 |
| BES4 | 431 | 13968 | 13538 | 913.47 | 50 bp repeats | 1.15 | 5.07 | 3.74 |
| BES4 | 51868 | 59054 | 7187 | 698.46 | 70 bp repeats upstream VSG gene | 2.87 | 127.69 | 125.17 |
| BES4 | 65997 | 68399 | 2403 | 622.09 | downstream VSG gene and telomeric repeats | 1.37 | 13.54 | 11.94 |
| BES5 | 20 | 1029 | 1010 | 874.64 | 50 bp repeats | 1.15 | 4.80 | 3.50 |
| BES5 | 7939 | 12700 | 4762 | 836.4 | 50 bp repeats | 1.29 | 12.61 | 10.90 |
| BES5 | 47686 | 61163 | 13478 | 731.7 | 70 bp repeats upstream VSG gene | 2.78 | 124.44 | 121.90 |
| BES5 | 64469 | 67357 | 2889 | 615.3 | downstream VSG gene and telomeric repeats | 1.42 | 17.41 | 15.71 |
| BES7 | 61 | 16589 | 16529 | 929.46 | 50 bp repeats | 1.20 | 7.97 | 6.41 |
| BES7 | 66175 | 80780 | 14606 | 727.95 | 70 bp repeats upstream VSG gene | 2.94 | 136.21 | 133.55 |
| BES7 | 85007 | 88058 | 3052 | 820.56 | downstream VSG gene and telomeric repeats | 1.47 | 26.19 | 24.36 |
| BES10 | 34 | 10778 | 10745 | 926.57 | 50 bp repeats | 1.27 | 12.30 | 10.58 |
| BES10 | 39600 | 41759 | 2160 | 709.85 | downstream VSG gene and telomeric repeats | 1.46 | 23.28 | 21.52 |
| BES11 | 15 | 11047 | 11033 | 966.74 | 50 bp repeats | 1.19 | 7.48 | 5.90 |
| BES11 | 45263 | 62863 | 17601 | 699.91 | 70 bp repeats upstream VSG gene | 2.53 | 101.66 | 99.36 |
| BES11 | 65577 | 66742 | 1166 | 568.65 | downstream VSG gene and telomeric repeats | 1.43 | 16.82 | 15.14 |
| BES12 | 56 | 5194 | 5139 | 923.27 | 50 bp repeats | 1.18 | 6.33 | 4.86 |
| BES12 | 37454 | 43618 | 6165 | 680.46 | 70 bp repeats upstream VSG gene | 2.72 | 114.64 | 112.27 |
| BES12 | 46973 | 47724 | 752 | 521.9 | downstream VSG gene and telomeric repeats | 1.46 | 16.99 | 15.25 |
| BES13 | 47 | 10393 | 10347 | 919.09 | 50 bp repeats | 1.18 | 6.45 | 4.97 |
| BES13 | 56050 | 60567 | 4518 | 667.55 | 70 bp repeats upstream VSG gene | 2.48 | 94.82 | 92.60 |
| BES14 | 35 | 6967 | 6933 | 905.02 | 50 bp repeats | 1.15 | 4.88 | 3.58 |
| BES14 | 39802 | 40843 | 1042 | 508.48 | 70 bp repeats upstream VSG gene | 2.12 | 57.68 | 55.74 |
| BES14 | 47089 | 77371 | 30283 | 724.28 | 70 bp repeats downstream VSG gene | 2.76 | 121.25 | 118.81 |
| BES15 | 19 | 14641 | 14623 | 946.04 | 50 bp repeats | 1.19 | 7.12 | 5.57 |
| BES15 | 60812 | 64074 | 3263 | 578.75 | 70 bp repeats downstream VSG gene | 2.41 | 85.69 | 83.44 |
| BES15 | 67372 | 85265 | 17894 | 705.71 | 70 bp repeats downstream VSG gene | 2.77 | 121.17 | 118.65 |
| BES15 | 90040 | 90926 | 887 | 400.45 | downstream VSG gene and telomeric repeats | 1.66 | 23.90 | 22.17 |
| BES17 | 33 | 11338 | 11306 | 926.79 | 50 bp repeats | 1.17 | 5.98 | 4.55 |
| BES17 | 56386 | 75949 | 19564 | 688.21 | 70 bp repeats downstream VSG gene | 2.80 | 120.72 | 118.32 |
| BES17 | 79092 | 86224 | 7133 | 737.25 | downstream VSG gene and telomeric repeats | 1.65 | 36.99 | 35.11 |
| MES_Chr10_5A | 1 | 1723 | 1723 | 855.27 | downstream VSG gene | 1.19 | 6.54 | 5.06 |
| MES_Chr10_5B | 3663 | 9535 | 5873 | 705.19 | upstream VSG gene | 2.85 | 127.74 | 125.13 |
| MES_Chr11_5A | 194 | 3564 | 3371 | 761.43 | downstream VSG gene and telomeric repeats | 1.40 | 18.92 | 17.19 |
| MES_Chr3_5A | 174 | 1010 | 837 | 518.61 | downstream VSG gene and telomeric repeats | 1.51 | 20.89 | 19.18 |
| MES_Chr8_5B | 61 | 1578 | 1518 | 842.2 | downstream VSG gene and telomeric repeats | 1.26 | 11.14 | 9.51 |
| MES_Chr9_5A | 1 | 723 | 723 | 483.81 | downstream VSG gene and telomeric repeats | 1.28 | 7.36 | 5.85 |
| MES_Chr9_5B | 10 | 607 | 598 | 509.81 | downstream VSG gene and telomeric repeats | 1.19 | 4.41 | 3.14 |
| Chr1_core | 602794 | 608865 | 6072 | 966.3 | centromere | 1.20 | 7.57 | 5.98 |
| Chr1_core | 610689 | 612940 | 2252 | 969.4 | centromere | 1.20 | 7.79 | 6.18 |
| Chr1_core | 614062 | 618411 | 4350 | 884.6 | centromere | 1.15 | 4.70 | 3.41 |
| Chr2_core | 16 | 8493 | 8478 | 940.74 | centromere | 1.19 | 7.19 | 5.63 |
| Chr3_core | 778606 | 802714 | 24109 | 926.76 | centromere | 1.18 | 6.43 | 4.94 |
| Chr4_core | 873852 | 879199 | 5348 | 981.22 | centromere | 1.19 | 7.50 | 5.92 |
| Chr4_core | 881038 | 882758 | 1721 | 964.26 | centromere | 1.18 | 6.50 | 5.00 |
| Chr5_core | 207144 | 223411 | 16268 | 985.49 | centromere | 1.20 | 7.66 | 6.07 |
| Chr5_core | 225239 | 228618 | 3380 | 973.39 | centromere | 1.19 | 7.42 | 5.85 |
| Chr6_core | 9 | 2672 | 2664 | 939.2 | centromere | 1.19 | 7.22 | 5.67 |
| Chr6_core | 3820 | 7920 | 4101 | 894.99 | centromere | 1.15 | 4.83 | 3.53 |
| Chr7_core | 1931029 | 1933296 | 2268 | 958.46 | centromere | 1.20 | 7.86 | 6.26 |
| Chr7_core | 1934404 | 1937614 | 3211 | 976.39 | centromere | 1.20 | 8.15 | 6.51 |
| Chr8_core | 2110966 | 2123617 | 12652 | 986 | centromere | 1.20 | 7.80 | 6.19 |
| Chr9_3A | 1342375 | 1345212 | 2838 | 947.54 | centromere | 1.20 | 7.64 | 6.05 |
| Chr9_3B | 287814 | 290363 | 2550 | 960.48 | centromere | 1.20 | 7.89 | 6.28 |
| Chr10_3A | 1249862 | 1256428 | 6567 | 971.76 | centromere | 1.20 | 7.73 | 6.13 |
| Chr10_3A | 1258244 | 1264378 | 6135 | 956.5 | centromere | 1.19 | 7.12 | 5.57 |
| Chr10_3B | 956393 | 967172 | 10780 | 970.96 | centromere | 1.20 | 8.03 | 6.40 |
| Chr11_3A | 774698 | 779063 | 4366 | 904.89 | centromere | 1.15 | 4.83 | 3.53 |
| Chr11_3A | 780410 | 784134 | 3725 | 995.63 | centromere | 1.21 | 8.43 | 6.77 |
| Chr11_3A | 785939 | 791468 | 5530 | 993.55 | centromere | 1.20 | 8.23 | 6.59 |
| Chr11_3A | 792782 | 797169 | 4388 | 885.86 | centromere | 1.15 | 4.78 | 3.49 |
| Chr11_3B | 901050 | 904797 | 3748 | 954.83 | centromere | 1.20 | 7.66 | 6.07 |
