## Supplementary File 5 for "PI(3,4,5)P3 allosteric regulation of repressor activator protein 1 controls antigenic variation in trypanosomes"

| **Experiment** | **Treatment** | **Replicates** | **Reads mean length** | **Sequences (bp)** |
| --- | --- | --- | --- | --- |
| RNA-seq | WT PIP5Pase | Bio1 | 668.919 | 2,072,167,984 |
|  |  | Bio2 | 2547.51 | 1,990,123,451 |
|  |  | Bio3 | 400.328 | 317,681,697 |
|  | Mut PIP5Pase | Bio1 | 639.227 | 1,697,997,595 |
|  |  | Bio2 | 492.199 | 320,814,615 |
|  |  | Bio3 | 237.398 | 533,192,072 |
| ChIP-seq | RAP1-HA in WT PIP5Pase | Bio1_Input | 323.63 | 465,606,176 |
|  |  | Bio1_ChIP | 369.573 | 588,132,165 |
|  |  | Bio2_Input | 722.822 | 879,935,255 |
|  |  | Bio2_ChIP | 587.867 | 257,141,699 |
|  |  | Bio3_Input | 749.543 | 698,570,934 |
|  |  | Bio3_ChIP | 285.618 | 678,626,652 |
|  | RAP1-HA in Mut PIP5Pase | Bio1_Input | 570.146 | 295,541,382 |
|  |  | Bio1_ChIP | 282.222 | 497,713,577 |
|  |  | Bio2_Input | 597.497 | 202,006,025 |
|  |  | Bio2_ChIP | 322.475 | 577,598,596 |
|  |  | Bio3_Input | 491.412 | 196,564,842 |
|  |  | Bio3_ChIP | 480.533 | 205,665,425 |
