## Supplementary File 6 for "PI(3,4,5)P3 allosteric regulation of repressor activator protein 1 controls antigenic variation in trypanosomes"

| **Oligonucleotide name** | **Sequence** |
| --- | --- |
| Tel-G-Biotin | 5’-/5Biosg/ TTAGGGTTAGGGTTAGGGTTAGGGTTAGGGTTAGGGTTAG |
|  | GG TTAGGGTTAGGGTTAGGG -3’ |
| Tel-G-RevCom | 5’-CCCTAACCCTAACCCTAACCCTAACCCTAACCCTAA CCCTACCCTA |
|  | ACCCTAACCCTAA -3’ |
| 70bp_Biotin | 5’-/5Biosg/ATGATAATGATAATAATAATAATAG -3’ |
| 70bp-RevCom | 5’-TATTATTACTGCTCTTATTATAATATTC -3’ |
| Tel-Scram-Biotin | 5’- /5Biosg/AGGTGGGTTGTGGTGTTGAGTTGTGTATAGATGGGGGAGG |
|  | TGGAGGTTAATGG -3’ |
| Tel-Scram-RevCom | 5’-ACAATCCCATTAACCTCCCACCTCCCCCATCTATACACAACTCAAC |
|  | ACCACCAACCCA -3’ |
| Tel-G-Cy5 | 5’-/5Cy5/TTAGGGTTAGGGTTAGGGTTAGGGTTAGGGTTAGGGTTAGG |
|  | GTTAGGGTTAGGGTTAGGG -3’ |
| 70bp_Cy5 | 5’-/5Cy5/ATGATAATGATAATAATAATAGGAGAGTGTTGTGAGAGTGT |
|  | ATATACGAATATT -3’ |
| Tel-Scram-Cy5 | 5’-/5Cy5/AGGTGGGTTGTGGTGTTGAGTTGTGTATAGATGGGGGAGGT |
|  | GGGAGGTTAATGG -3’ |
| SaCas9g_RAP1-76 | 5’-ACTCACTATAGGGAGACCACTTCGCGAAGTTTATACATAGGATGT |
|  | TTTAGTACTCTGT -3’ |
| SaCas9g_RAP1-2708 | 5’-ACTCACTATAGGGAGAGGTGCGTCTGTTGAGATCACAAAGAGTGT |
|  | TTTAGTACTCTGT -3’ |
| rRAP1^1-300^-F | 5’- CCCCATATGACCGACGTTGACGGT -3’ |
| rRAP1^1-300^-R | 5’- CCCCTCGAGGTCATCGCTTTCATCAACTTCG -3’ |
| rRAP1^301-560^-F | 5’- CCCCATATGAGCATAAAGGACGAGTTCAGTTACAG -3’ |
| rRAP1^301-560^-R | 5’- CCCCTCGAGCGATTCCTTGGAACCTTCATT -3’ |
| rRAP1^561-855^-F | 5’- CCCCATATGGCTACCCCTGCTCTTCCTTTTC -3’ |
| rRAP1^561-855^-R | 5’- CCCCTCGAGTTGTGGCGCTTTCGAAGT -3’ |
| VSG_Splice Leader | 5’-ACAGTTTCTGTACTATATTG -3’ |
| SP6-VSG14mer | 5’- GATTTAGGTGACACTATAGTGTTAAAATATATC -3’ |
| BCA_SL | 5’- TTTCTGTTGGTGCTGATATTGCACAGTTTCTGTACTATATTG -3’ |
| BCA_Rd_3'AllVSGs_1 | 5’-TACTTGCCTGTCGCTCTATCTTCCGTTAGGTGTTAAAATATATC -3’ |
| BCA_Rd_3'AllVSGs_2 | 5’-TACTTGCCTGTCGCTCTATCTTCCAACTAGTGTTAAAATATATC -3’ |
| BCA_Rd_3'AllVSGs_3 | 5’-TACTTGCCTGTCGCTCTATCTTCAGGGCAGTGTTAAAATATATC -3’ |
| BCA_Rd_3'AllVSGs_4 | 5’-TACTTGCCTGTCGCTCTATCTTCGCAGAAGTGTTAAAATATATC -3’ |
| BCA_Rd_3'AllVSGs_5 | 5’-TACTTGCCTGTCGCTCTATCTTCGCTAAAGTGTTAAAATATATC -3’ |
| BCA_Rd_3'AllVSGs_6 | 5’-TACTTGCCTGTCGCTCTATCTTCCGCCCTGTGTTAAAATATATC -3’ |
| BCA_Rd_3'AllVSGs_7 | 5’-TACTTGCCTGTCGCTCTATCTTCTCCAAAGTGTTAAAATATATC -3’ |
| BCA_Rd_3'AllVSGs_8 | 5’-TACTTGCCTGTCGCTCTATCTTCATTACCGTGTTAAAATATATC -3’ |
| BCA_Rd_3'AllVSGs_9 | 5’-TACTTGCCTGTCGCTCTATCTTCCCGAACGTGTTAAAATATATC -3’ |
| BCA_Rd_3'AllVSGs_10 | 5’-TACTTGCCTGTCGCTCTATCTTCACACTCGTGTTAAAATATATC -3’ |
| BCA_Rd_3'AllVSGs_11 | 5’-TACTTGCCTGTCGCTCTATCTTCCAACAAGTGTTAAAATATATC -3’ |
| BCA_Rd_3'AllVSGs_12 | 5’-TACTTGCCTGTCGCTCTATCTTCGAACTCGTGTTAAAATATATC -3’ |
| BCA_Rd_3'AllVSGs_13 | 5’-TACTTGCCTGTCGCTCTATCTTCCGAAATGTGTTAAAATATATC -3’ |
| BCA_Rd_3'AllVSGs_14 | 5’-TACTTGCCTGTCGCTCTATCTTCATCGCAGTGTTAAAATATATC -3’ |
| BCA_Rd_3'AllVSGs_15 | 5’-TACTTGCCTGTCGCTCTATCTTCATTTTAGTGTTAAAATATATC -3’ |
| BCA_Rd_3'AllVSGs_16 | 5’-TACTTGCCTGTCGCTCTATCTTCTCGCAAGTGTTAAAATATATC -3’ |
| BCA_Rd_3'AllVSGs_17 | 5’-TACTTGCCTGTCGCTCTATCTTCCGGCATGTGTTAAAATATATC -3’ |
| BCA_Rd_3'AllVSGs_18 | 5’-TACTTGCCTGTCGCTCTATCTTCGGTCGAGTGTTAAAATATATC -3’ |
| BCA_Rd_3'AllVSGs_19 | 5’-TACTTGCCTGTCGCTCTATCTTCTATAAAGTGTTAAAATATATC -3’ |
| BCA_Rd_3'AllVSGs_20 | 5’-TACTTGCCTGTCGCTCTATCTTCAACTATGTGTTAAAATATATC -3’ |
| BCA_Rd_3'AllVSGs_21 | 5’-TACTTGCCTGTCGCTCTATCTTCGACAATGTGTTAAAATATATC -3’ |
| BCA_Rd_3'AllVSGs_22 | 5’-TACTTGCCTGTCGCTCTATCTTCGAAACGGTGTTAAAATATATC -3’ |
| BCA_Rd_3'AllVSGs_23 | 5’-TACTTGCCTGTCGCTCTATCTTCGTTCCCGTGTTAAAATATATC -3’ |
| BCA_Rd_3'AllVSGs_24 | 5’-TACTTGCCTGTCGCTCTATCTTCACGGATGTGTTAAAATATATC -3’ |
| BCA_Rd_3'AllVSGs_25 | 5’-TACTTGCCTGTCGCTCTATCTTCACGAAAGTGTTAAAATATATC -3’ |
| BCA_Rd_3'AllVSGs_26 | 5’-TACTTGCCTGTCGCTCTATCTTCTAAGGGGTGTTAAAATATATC -3’ |
| BCA_Rd_3'AllVSGs_27 | 5’-TACTTGCCTGTCGCTCTATCTTCACCTCGGTGTTAAAATATATC -3’ |
| BCA_Rd_3'AllVSGs_28 | 5’-TACTTGCCTGTCGCTCTATCTTCAAACGCGTGTTAAAATATATC -3’ |
| BCA_Rd_3'AllVSGs_29 | 5’-TACTTGCCTGTCGCTCTATCTTCGTCGTAGTGTTAAAATATATC -3’ |
| BCA_Rd_3'AllVSGs_30 | 5’-TACTTGCCTGTCGCTCTATCTTCCGTAGGGTGTTAAAATATATC -3’ |
| BCA_Rd_3'AllVSGs_31 | 5’-TACTTGCCTGTCGCTCTATCTTCTAAGCGGTGTTAAAATATATC -3’ |
| BCA_Rd_3'AllVSGs_32 | 5’-TACTTGCCTGTCGCTCTATCTTCTATGGAGTGTTAAAATATATC -3’ |
| BCA_Rd_3'AllVSGs_33 | 5’-TACTTGCCTGTCGCTCTATCTTCCGGCCTGTGTTAAAATATATC -3’ |
| BCA_Rd_3'AllVSGs_34 | 5’-TACTTGCCTGTCGCTCTATCTTCCCGGGCGTGTTAAAATATATC -3’ |
| BCA_Rd_3'AllVSGs_35 | 5’-TACTTGCCTGTCGCTCTATCTTCTGGTCGGTGTTAAAATATATC -3’ |
| BCA_Rd_3'AllVSGs_36 | 5’-TACTTGCCTGTCGCTCTATCTTCGTAACGGTGTTAAAATATATC -3’ |
| BCA_Rd_3'AllVSGs_37 | 5’-TACTTGCCTGTCGCTCTATCTTCGGTACGGTGTTAAAATATATC -3’ |
| BCA_Rd_3'AllVSGs_38 | 5’-TACTTGCCTGTCGCTCTATCTTCCTGAGTGTGTTAAAATATATC -3’ |
| BCA_Rd_3'AllVSGs_39 | 5’-TACTTGCCTGTCGCTCTATCTTCTACGCGGTGTTAAAATATATC -3’ |
| BCA_Rd_3'AllVSGs_40 | 5’-TACTTGCCTGTCGCTCTATCTTCAATTTGGTGTTAAAATATATC -3’ |
| BCA_Rd_3'AllVSGs_41 | 5’-TACTTGCCTGTCGCTCTATCTTCGTGCCCGTGTTAAAATATATC -3’ |
| BCA_Rd_3'AllVSGs_42 | 5’-TACTTGCCTGTCGCTCTATCTTCCCTACTGTGTTAAAATATATC -3’ |
| BCA_Rd_3'AllVSGs_43 | 5’-TACTTGCCTGTCGCTCTATCTTCATACCGGTGTTAAAATATATC -3’ |
| BCA_Rd_3'AllVSGs_44 | 5’-TACTTGCCTGTCGCTCTATCTTCGCGGAGGTGTTAAAATATATC -3’ |
| BCA_Rd_3'AllVSGs_45 | 5’-TACTTGCCTGTCGCTCTATCTTCGAACCTGTGTTAAAATATATC -3’ |
| BCA_Rd_3'AllVSGs_46 | 5’-TACTTGCCTGTCGCTCTATCTTCTTAATTGTGTTAAAATATATC -3’ |
| BCA_Rd_3'AllVSGs_47 | 5’-TACTTGCCTGTCGCTCTATCTTCGTATAGGTGTTAAAATATATC -3’ |
| BCA_Rd_3'AllVSGs_48 | 5’-TACTTGCCTGTCGCTCTATCTTCGTCAAGGTGTTAAAATATATC -3’ |
| BCA_Rd_3'AllVSGs_49 | 5’-TACTTGCCTGTCGCTCTATCTTCTCGCACGTGTTAAAATATATC -3’ |
| BCA_Rd_3'AllVSGs_50 | 5’-TACTTGCCTGTCGCTCTATCTTCGTCTGCGTGTTAAAATATATC -3’ |
| BCA_Rd_3'AllVSGs_51 | 5’-TACTTGCCTGTCGCTCTATCTTCAGTAAGGTGTTAAAATATATC -3’ |
| BCA_Rd_3'AllVSGs_52 | 5’-TACTTGCCTGTCGCTCTATCTTCGTTCCCGTGTTAAAATATATC -3’ |
| BCA_Rd_3'AllVSGs_53 | 5’-TACTTGCCTGTCGCTCTATCTTCTTGCATGTGTTAAAATATATC -3’ |
| BCA_Rd_3'AllVSGs_54 | 5’-TACTTGCCTGTCGCTCTATCTTCTAAAATGTGTTAAAATATATC -3’ |
| BCA_Rd_3'AllVSGs_55 | 5’-TACTTGCCTGTCGCTCTATCTTCTCTCATGTGTTAAAATATATC -3’ |
| BCA_Rd_3'AllVSGs_56 | 5’-TACTTGCCTGTCGCTCTATCTTCGGTTGGGTGTTAAAATATATC -3’ |
| BCA_Rd_3'AllVSGs_57 | 5’-TACTTGCCTGTCGCTCTATCTTCGCACGTGTGTTAAAATATATC -3’ |
| BCA_Rd_3'AllVSGs_58 | 5’-TACTTGCCTGTCGCTCTATCTTCGTCAAAGTGTTAAAATATATC -3’ |
| BCA_Rd_3'AllVSGs_59 | 5’-TACTTGCCTGTCGCTCTATCTTCCGTGAAGTGTTAAAATATATC -3’ |
| BCA_Rd_3'AllVSGs_60 | 5’-TACTTGCCTGTCGCTCTATCTTCGACACCGTGTTAAAATATATC -3’ |
| BCA_Rd_3'AllVSGs_61 | 5’-TACTTGCCTGTCGCTCTATCTTCCCGCCAGTGTTAAAATATATC -3’ |
| BCA_Rd_3'AllVSGs_62 | 5’-TACTTGCCTGTCGCTCTATCTTCGGACAAGTGTTAAAATATATC -3’ |
| BCA_Rd_3'AllVSGs_63 | 5’-TACTTGCCTGTCGCTCTATCTTCTATGCGGTGTTAAAATATATC -3’ |
| BCA_Rd_3'AllVSGs_64 | 5’-TACTTGCCTGTCGCTCTATCTTCGGCGTTGTGTTAAAATATATC -3’ |
| BCA_Rd_3'AllVSGs_65 | 5’-TACTTGCCTGTCGCTCTATCTTCACGAATGTGTTAAAATATATC -3’ |
| BCA_Rd_3'AllVSGs_66 | 5’-TACTTGCCTGTCGCTCTATCTTCACCGGGGTGTTAAAATATATC -3’ |
| BCA_Rd_3'AllVSGs_67 | 5’-TACTTGCCTGTCGCTCTATCTTCCGAGGGGTGTTAAAATATATC -3’ |
| BCA_Rd_3'AllVSGs_68 | 5’-TACTTGCCTGTCGCTCTATCTTCCGGCATGTGTTAAAATATATC -3’ |
| BCA_Rd_3'AllVSGs_69 | 5’-TACTTGCCTGTCGCTCTATCTTCAAGTCGGTGTTAAAATATATC -3’ |
| BCA_Rd_3'AllVSGs_70 | 5’-TACTTGCCTGTCGCTCTATCTTCATGCGTGTGTTAAAATATATC -3’ |
| BCA_Rd_3'AllVSGs_71 | 5’-TACTTGCCTGTCGCTCTATCTTCAGCGATGTGTTAAAATATATC -3’ |
| BCA_Rd_3'AllVSGs_72 | 5’-TACTTGCCTGTCGCTCTATCTTCTAATGTGTGTTAAAATATATC -3’ |
| BCA_Rd_3'AllVSGs_73 | 5’-TACTTGCCTGTCGCTCTATCTTCCCGCCTGTGTTAAAATATATC -3’ |
| BCA_Rd_3'AllVSGs_74 | 5’-TACTTGCCTGTCGCTCTATCTTCCGTACCGTGTTAAAATATATC -3’ |
| BCA_Rd_3'AllVSGs_75 | 5’-TACTTGCCTGTCGCTCTATCTTCGCCGTCGTGTTAAAATATATC -3’ |
| BCA_Rd_3'AllVSGs_76 | 5’-TACTTGCCTGTCGCTCTATCTTCACCGCGGTGTTAAAATATATC -3’ |
| BCA_Rd_3'AllVSGs_77 | 5’-TACTTGCCTGTCGCTCTATCTTCCATGGGGTGTTAAAATATATC -3’ |
| BCA_Rd_3'AllVSGs_78 | 5’-TACTTGCCTGTCGCTCTATCTTCCTAGTTGTGTTAAAATATATC -3’ |
| BCA_Rd_3'AllVSGs_79 | 5’-TACTTGCCTGTCGCTCTATCTTCGTGATAGTGTTAAAATATATC -3’ |
| BCA_Rd_3'AllVSGs_80 | 5’-TACTTGCCTGTCGCTCTATCTTCTTGGACGTGTTAAAATATATC -3’ |
| BCA_Rd_3'AllVSGs_81 | 5’-TACTTGCCTGTCGCTCTATCTTCAGTATCGTGTTAAAATATATC -3’ |
| BCA_Rd_3'AllVSGs_82 | 5’-TACTTGCCTGTCGCTCTATCTTCTTCGATGTGTTAAAATATATC -3’ |
| BCA_Rd_3'AllVSGs_83 | 5’-TACTTGCCTGTCGCTCTATCTTCCTCGACGTGTTAAAATATATC -3’ |
| BCA_Rd_3'AllVSGs_84 | 5’-TACTTGCCTGTCGCTCTATCTTCTCGCTAGTGTTAAAATATATC -3’ |
| BCA_Rd_3'AllVSGs_85 | 5’-TACTTGCCTGTCGCTCTATCTTCAGTTGGGTGTTAAAATATATC -3’ |
| BCA_Rd_3'AllVSGs_86 | 5’-TACTTGCCTGTCGCTCTATCTTCCGGGGTGTGTTAAAATATATC -3’ |
| BCA_Rd_3'AllVSGs_87 | 5’-TACTTGCCTGTCGCTCTATCTTCCATCCCGTGTTAAAATATATC -3’ |
| BCA_Rd_3'AllVSGs_88 | 5’-TACTTGCCTGTCGCTCTATCTTCTAATGCGTGTTAAAATATATC -3’ |
| BCA_Rd_3'AllVSGs_89 | 5’-TACTTGCCTGTCGCTCTATCTTCATGGCGGTGTTAAAATATATC -3’ |
| BCA_Rd_3'AllVSGs_90 | 5’-TACTTGCCTGTCGCTCTATCTTCGGGTTTGTGTTAAAATATATC -3’ |
| BCA_Rd_3'AllVSGs_91 | 5’-TACTTGCCTGTCGCTCTATCTTCAGCTACGTGTTAAAATATATC -3’ |
| BCA_Rd_3'AllVSGs_92 | 5’-TACTTGCCTGTCGCTCTATCTTCGCGAATGTGTTAAAATATATC -3’ |
| BCA_Rd_3'AllVSGs_93 | 5’-TACTTGCCTGTCGCTCTATCTTCTAAACGGTGTTAAAATATATC -3’ |
| BCA_Rd_3'AllVSGs_94 | 5’-TACTTGCCTGTCGCTCTATCTTCTAATGAGTGTTAAAATATATC -3’ |
| BCA_Rd_3'AllVSGs_95 | 5’-TACTTGCCTGTCGCTCTATCTTCGTACCCGTGTTAAAATATATC -3’ |
| BCA_Rd_3'AllVSGs_96 | 5’-TACTTGCCTGTCGCTCTATCTTCCCAGACGTGTTAAAATATATC -3’ |
